# The L1 stalk is required for efficient export of nascent large ribosomal subunits in yeast

**DOI:** 10.1101/629261

**Authors:** Sharmishtha Musalgaonkar, Joshua J. Black, Arlen W. Johnson

## Abstract

The ribosomal protein Rpl1 (uL1 in universal nomenclature) is essential in yeast and constitutes part of the L1 stalk which interacts with E site ligands on the ribosome. Structural studies of nascent pre-60S complexes in yeast have shown that a domain of the Crm1-dependent nuclear export adapter Nmd3, binds in the E site and interacts with Rpl1, inducing closure of the L1 stalk. Based on this observation, we decided to reinvestigate the role of the L1 stalk in nuclear export of pre-60S subunits despite previous work showing that Rpl1-deficient ribosomes are exported from the nucleus and engage in translation. Large cargoes, such as ribosomal subunits, require multiple export factors to facilitate their transport through the nuclear pore complex. Here, we show that pre-60S subunits lacking Rpl1 or truncated for the RNA of the L1 stalk are exported inefficiently. Surprisingly, this is not due to a measurable defect in recruitment of Nmd3 but appears to result from inefficient recruitment of the Mex67-Mtr2 heterodimer.

## Introduction

Ribosomes are universally composed of one large and one small subunit, that function together to synthesize all proteins in a cell. Production of balanced levels of ribosomal subunits is critical for maintaining homeostasis in cells. In yeast four rRNA molecules and about 80 ribosomal proteins interact with more than 200 trans-acting assembly factors to achieve the complex task of ribosome synthesis (Woolford and Baserga 2013). Synthesis of new ribosomes by cells is a challenging and energy consuming task and requires the coordinated expression of all ribosomal proteins and rRNAs. In yeast, failure to establish balanced expression levels of ribosomal proteins has been reported to cause cellular stress (Boulon et al. 2010; Cheng et al. 2019). Haploinsufficiency and mutations in ribosomal proteins in drosophila and zebrafish cause defects and delays in development (Amsterdam et al. 2004). In humans, mutations in genes coding for ribosomal proteins and biogenesis factors, give rise to a special class of diseases called ribosomopathies (De Keersmaecker et al. 2013; Mills and Green 2017; Warren 2018).

In eukaryotic cells, ribosome assembly starts in the sub-nuclear compartment called the nucleolus; it continues in the nucleoplasm followed by nuclear export; and concludes in the cytoplasm rendering fully matured subunits (reviewed in(Woolford and Baserga 2013; Kressler et al. 2017; Peña et al. 2017)). Although, the precursor rRNA for both subunits is a single transcript, RNA processing in the nucleolus separates precursors for the two subunits before nuclear export. Much of ribosomal subunit assembly is completed in the nucleus before the subunits are exported to the cytoplasm. The nuclear export machinery has to therefore undertake a critical task of escorting the highly hydrophilic and bulky pre-ribosomal subunits through hydrophobic environment of the nuclear pore complex (NPC). It was previously shown that large cargoes require multiple receptor molecules for effecting transient and reversible collapse of the hydrophobic permeability barrier in NPC for rapid translocation (Ribbeck and Görlich 2002). Consistent with this model, several export factors are required for the export of nascent 60S subunits. Export is facilitated by the export adaptor protein Nmd3 that utilizes its nuclear export sequence (NES) to recruit the export-receptor Crm1 (Ho et al. 2000; Gadal et al. 2001). Nuclear export is also assisted by other non-canonical export factors including Arx1 (Hung et al. 2007; Greber et al. 2012, 2016; Bradatsch et al. 2007) and Bud20 (Bassler et al. 2012; Altvater et al. 2012) and the mRNA export factor Mex67-Mtr2 heterodimer (Yao et al. 2007) in yeast but only Nmd3 appears to be conserved throughout eukaryotes as a dedicated 60S export factor (Thomas and Kutay 2003). However, unlike its essential role in 60S biogenesis, Nmd3 interaction with Crm1 is dispensable if other export receptors are fused directly to Nmd3 (Lo and Johnson 2009) indicating a general requirement for interaction with the NPC but not a specific requirement for a particular export receptor.

Recent high resolution structures of ribosome assembly intermediates have revealed the binding sites of numerous biogenesis factors including all known export factors with the exception of Mex67-Mtr2 (Zhou et al. 2019; Malyutin et al. 2017; Greber et al. 2016; Wu et al. 2016; Ma et al. 2017; Matsuo et al. 2014; Barrio-Garcia et al. 2016). Arx1 binds on the solvent-exposed surface of the subunit near the peptide exit tunnel (PET) whereas Nmd3 binds on the subunit interface spanning the E, P and A-sites of the subunit. However, the C-terminal region of Nmd3 which contains the NES that recruits Crm1, is not resolved on any of the structures. Therefore, the position of Crm1 relative to the subunit during export is still unknown. Bud20 also binds on the subunit interface where it interacts with Tif6 and Rlp24, although the mechanism by which Bud20 promotes export is disputed (Bassler et al. 2012; Altvater et al. 2012). Densities for Mex67-Mtr2 heterodimer were not detected in any structures of pre-60S particles to date. However, UV-induced protein-RNA crosslinking studies *in vivo* identified crosslinks to many regions of the 25S rRNA, but strongly enriched in the 3’ terminal end of 5.8S rRNA (Tuck and Tollervey 2013). In addition, a recent study attempting to reconstitute Mex67-Mtr2 binding to affinity-purified pre-60S particles *in vitro* identified crosslinks to 5.8S and P-stalk rRNA (Sarkar et al. 2016). However, binding to the P-stalk was only observed for Yvh1-containing pre-60S particles while our recent structural studies and work from others show that Yvh1 joins the subunit only in the cytoplasm (Kemmler et al. 2009; Lo et al. 2009; Nerurkar et al. 2018; Zhou et al. 2019) and hence, cannot be responsible for promoting Mex67-Mtr2 binding to pre-60S in the nucleus.

Ribosomal Protein L1 (Rpl1) is universally conserved protein which interacts with a single loop of the 25S rRNA to form the L1 stalk. Rpl1 is essential in yeast and is encoded by two paralogous genes *RPL1A* and *RPL1B*. It was previously reported that 60S subunits lacking Rpl1 are exported to the cytoplasm and even detected in the polysomes (McIntosh et al. 2011), suggesting that 60S assembly and export can proceed in the absence of Rpl1. However, the essential export adapter Nmd3 binds Rpl1 and facilitates closure of the L1 stalk (Malyutin et al. 2017; Zhou et al. 2019; Ma et al. 2017). Because of the interaction between Rpl1 and Nmd3, we suspected that ribosomes lacking Rpl1 would be affected in their ability to bind Nmd3 or to release it.

Here we show that Rpl1 protein is needed for efficient nuclear export of nascent large subunits precursors. The repression of the *RPL1* or truncation of the L1 stalk rRNA reduced the efficiency of export but did not completely block export from the nucleus. Nascent subunits lacking Rpl1 maintained binding to the export factors Nmd3, Arx1 and Bud20 but only inefficiently recruited the Mex67-Mtr2 heterodimer. Co-overexpression of *MEX67* and *MTR2* in *RPL1* repressed cells overcame the 60S export defect caused by loss of Rpl1 suggesting that delayed export of Rpl1-deficient subunits is due to a failure in Mex67-Mtr2 recruitment.

## Results

### Repression of Rpl1 leads to a 60S subunit export defect

Recent cryo-EM structures of the nuclear export adapter Nmd3 on the 60S subunit revealed a large interface between the eIF5A-like domain of Nmd3 and ribosomal protein Rpl1 on the L1 stalk (Malyutin et al. 2017; Zhou et al. 2019; Ma et al. 2017). The interaction between Nmd3 and Rpl1 holds the L1 stalk in a closed conformation in which the L1 stalk is bent toward the E site. Conceivably, the interaction between Rpl1 and Nmd3 may be important for the recruitment of Nmd3 to the pre-60S subunit in the nucleus. Alternatively, this compact structure could facilitate export of the nascent 60S subunit through the nuclear pore complex or facilitate the release of Nmd3 from the pre-60S subunit after export to the cytoplasm. Although previous work reported that Rpl1 was not needed for 60S export (McIntosh et al. 2011), we decided to revisit this question. We first asked if nuclear export of 60S subunits was affected by loss of Rpl1. In yeast, Rpl1 is expressed from two paralogous genes, *RPL1A* and *RPL1B*, each encoding identical proteins. Because deletion of both genes is lethal, we used a conditional mutant in which *RPL1B* was deleted and *RPL1A* was under control of the galactose inducible/glucose repressible *GAL1* promoter. 60S subunit localization was monitored with a GFP fusion to Rpl25. Upon shifting cells from galactose to glucose to repress RPl1A expression, we observed a strong change in Rpl25-GFP localization from the cytoplasm to the nucleus (Fig. 1A), indicating impaired 60S export. Similar results were obtained using Rpl8-GFP as a reporter (data not shown). The accumulation was most evident within 60 to 90 minutes after returning saturated cultures to active growth. At longer time points, Rpl25 signal became increasingly cytoplasmic, indicating that export continued, albeit at a slower rate than when RPl1A was expressed (data not shown). Nuclear accumulation of Rpl25-GFP was also observed in an *rpl1b*∆ strain in which Rpl1 was constitutively expressed at reduced levels compared to wild-type cells due to deletion of *RPL1B* (see below).

**Figure 1.**
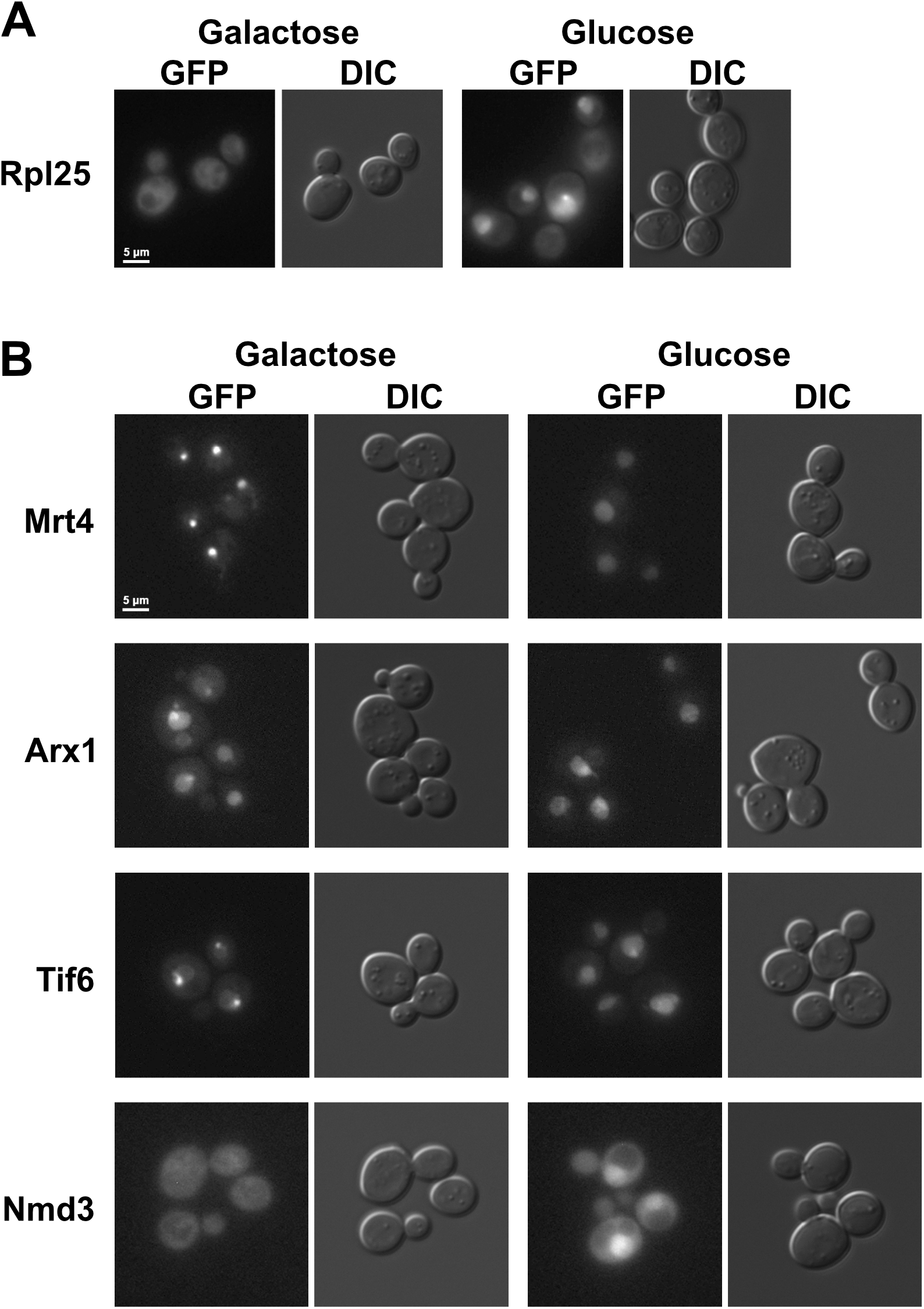
Depletion of Rpl1 reduces 60S export. A) The localization of Rpl25-GFP expressed from plasmid pAJ907 was visualized in cells of strain KBM20 expressing *RPL1A* (Galactose) or after 2 hours of repression of *RPL1A* (Glucose). GFP, tagged Rpl25; DIC, differential interference contrast. B) The localization of Mrt4-GFP (AJY3850), Tif6-GFP (AJY3848), Arx1-GFP (AJY3849) and Nmd3-GFP (AJY4060) was visualized in cells expressing *RPL1A* (Galactose) or after 2 hrs of repression of *RPL1A* (Glucose).

Pre-60S subunits are accompanied to the cytoplasm with a host of assembly factors including Nmd3, Mrt4, Tif6 and Arx1(Zhou et al. 2019; Ma et al. 2017; Wu et al. 2016; Barrio-Garcia et al. 2016). Although each of these factors continually shuttles in and out of the nucleus, they show different steady state localizations: Mrt4 and Tif6 are predominantly nucleolar whereas Arx1 is nucleoplasmic and Nmd3 is cytoplasmic (Lo et al. 2010). Upon glucose repression of *RPL1* expression, Mrt4 and Tif6 re-localized from the nucleolus to the nucleoplasm whereas the nucleoplasmic localization of Arx1 was largely unchanged (Fig. 1B). Strikingly, Nmd3 was relocalized from the cytoplasm to the nucleoplasm (Fig. 1B). These results imply that in the absence of Rpl1 expression, pre-60S particles containing Tif6, Mrt4, and probably Arx1 and Nmd3 accumulate at a late assembly step in the nucleoplasm, prior to export. We conclude that Rpl1 is necessary for efficient export of 60S subunits from the nucleus.

### Nmd3 binds to nascent subunits lacking Rpl1

Because Nmd3 binds to Rpl1 and provides an essential Crm1-dependent nuclear export signal for the 60S subunit, a lack of Nmd3 binding to the pre-60S particle could explain the export block observed upon repression of Rpl1. We tested if Nmd3 is lost from nascent 60S subunits under conditions where we observed accumulation of Nmd3 in the nucleus after Rpl1 repression. Cells expressing Rpl1 from its native promoters or under control of the glucose-repressible *GAL1* promoter were shifted from galactose to glucose. Extracts were prepared and sedimented through sucrose density gradients and the positions of Nmd3, Rpl1 and Rpl8 were monitored by western blotting. Surprisingly, Nmd3 co-sedimentation with free 60S subunits was largely unaffected by *RPL1* repression (Fig. 2, compare panel B with A). The slight accumulation of free Nmd3 at the top of the gradient in Rpl1-repressed cells (Fig. 2B) cannot account for the bulk redistribution of Nmd3 from the cytoplasm to the nucleus in these cells. These results suggest that the population of Nmd3 that accumulates in the nucleus upon *RPL1* repression is bound to pre-60S subunits.

**Figure 2.**
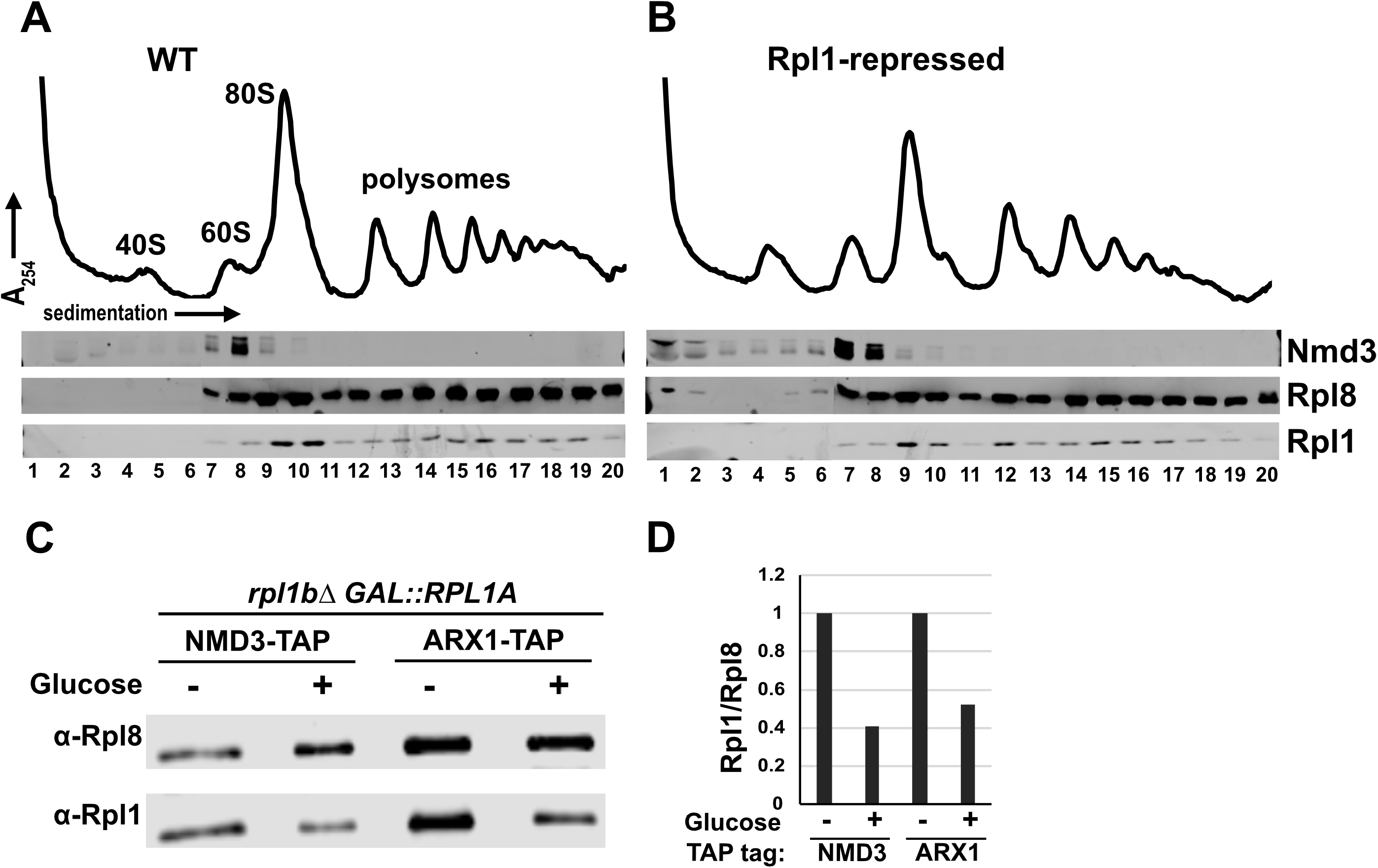
Nmd3 binds to subunits lacking Rpl1. A and B) Polysome profiles and western blots for monitoring sedimentation of Nmd3, Rpl1 and Rpl8 from extracts of WT (KBM13) and *GAL:: RPL1* (KBM20) cells, respectively, grown in galactose media followed by 2h growth after adding glucose. C) Western blots for affinity purification of Nmd3-TAP (AJY4009, lanes 1and 2) and Arx1-TAP (AJY4013, lanes 3 and 4) from *GAL::RPL1* cells either grown in galactose medium continually (lanes 1 and 3) or for 2h after addition of glucose (lanes 2 and 4). D) Ratios of Rpl1 to Rpl8 signal from western blot in C were calculated and normalized to the Rpl1 to Rpl8 ratios for cells grown continuously in galactose medium.

To test directly if Nmd3 binds to subunits lacking Rpl1, we immunoprecipitated subunits associated with Nmd3 from cells in which Rpl1 was expressed or repressed. We used Arx1 as a control for a pre-60S associated protein whose binding is not expected to be dependent on Rpl1. Similar levels of 60S subunits, indicated by Rpl8, coimmunoprecipitated with Nmd3 and Arx1 regardless of Rpl1 expression (Fig. 2C). However, the ratio of Rpl1 to Rpl8 in the immunoprecipitated samples was significantly reduced by when Rpl1 was repressed (Fig. 2D). These results show that Nmd3 can bind subunits lacking Rpl1, despite the loss of a large interaction surface between these two proteins.

### Truncation of the L1 stalk leads to a 60S export defect

As a complementary means of assessing the importance of the L1 stalk for 60S export, we truncated the RNA of the L1 stalk. We replaced nucleotides 2451-2495 with the GNRA tetraloop GAGA (Ben Shem et al. 2011; Correll et al. 1999), deleting the entire Rpl1 binding site (Fig. 3A). We made the truncation in a construct that ectopically expressed 25S rRNA with a unique oligo tag to be able to monitor the mutant ribosomal RNA in the presence of wild-type 60S. The oligo tag was inserted in ES8 and had no discernible effect on function (Fig. 3B). As anticipated, truncation of the L1 stalk was lethal, shown by its inability to complement deletion of the genomic rDNA locus (Fig. 3B). Nevertheless, we were able to monitor localization and incorporation of the mutant rRNA into subunits using fluorescence in situ hybridization (FISH) and northern blotting, respectively. The RNA of the L1 stalk truncation mutant accumulated in the nucleus but could also be detected in the cytoplasm (Fig. 3C), indicating that subunits with a truncated L1 stalk, and hence lacking Rpl1, can be exported to the cytoplasm, albeit less efficiently than wild-type. Surprisingly, this RNA sedimented in 60S, 80S and in polysomes, indicating that despite lacking a functional L1 stalk, the mutant RNA was incorporated into actively translating ribosomes. This was consistent with a previous report that Rpl1-deficient ribosomes can engage in translation (McIntosh et al. 2011). However, comparison of the ratio 25S rRNA from the L1 stalk∆ mutant to endogenously expressed 25S rRNA revealed differences in sedimentation of the mutant RNA compared to WT. Notably, the mutant strongly accumulated in free 60S subunits (Fig 3D, right panel, lanes 6 and 7) and was somewhat enriched over wild-type in 80S and light polysomes (fractions 10-15) but was relatively depleted from deep polysomes (fractions 16-19). A shift towards lighter polysomes suggests that ribosomes without a functional L1 stalk arrest at or shortly after translation initiation. Together, these results indicate that ribosomes with a truncated L1 stalk are exported slowly and engage with 40S subunits but accumulate in light polysomes, possibly because they are defective for elongation.

**Figure 3.**
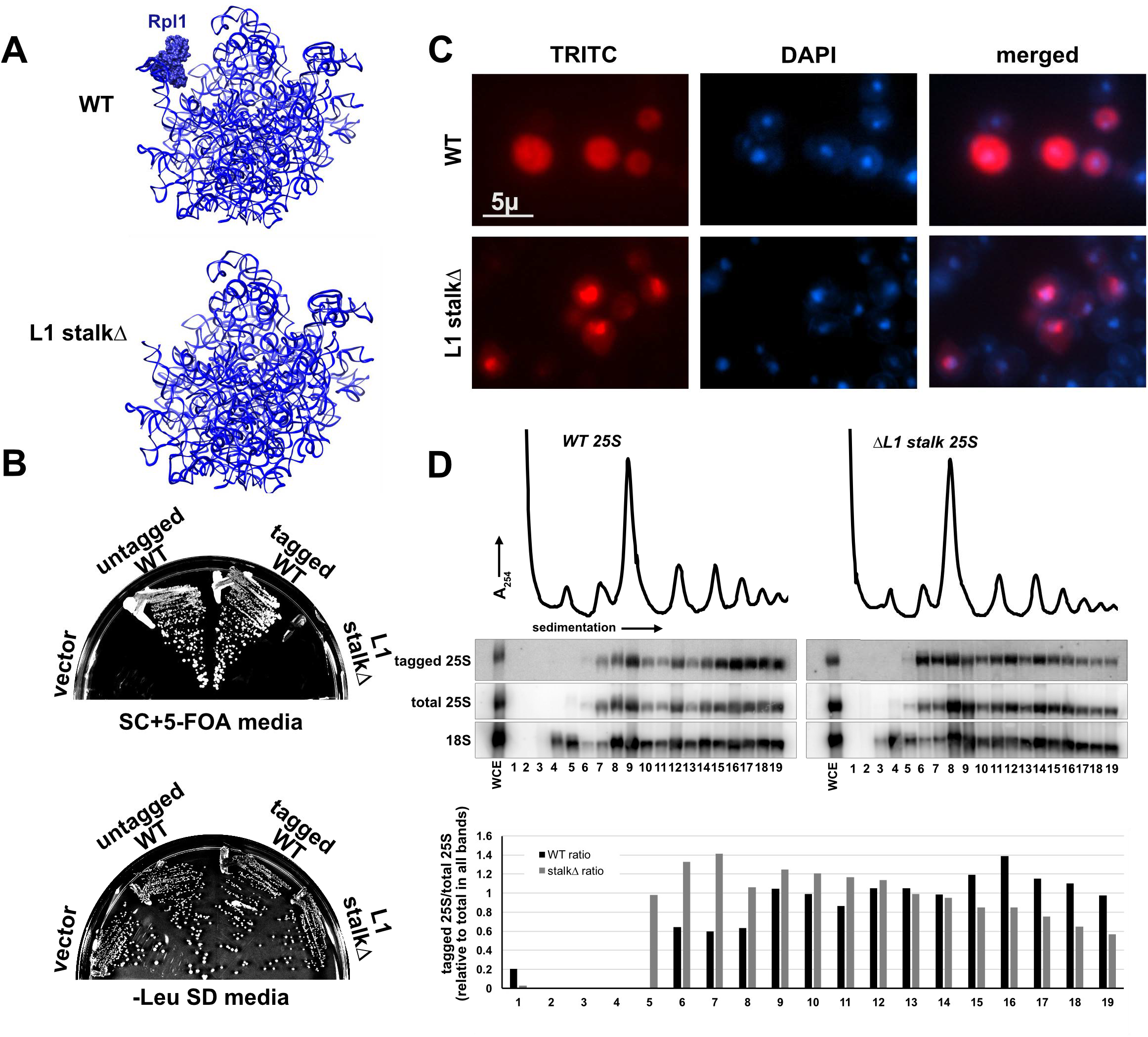
Truncation of L1 stalk RNA leads to a 60S export defect. A) Cartoon of 25S rRNA for WT and L1-stalk truncation showing expected lack of Rpl1 binding when RNA was truncated. B) Plasmids constructs expressing WT (pAJ1181) or L1 stalk∆ (pAJ3605) rRNA were transformed into AJY1185 (rDNA∆, 35S URA3 2µ) and complementation was tested on 5-FOA media. C) Fluorescence *in situ* hybridization and microscopy using oligo (AJO628) hybridizing to a unique tag in 25S rRNA expressed from plasmid constructs for WT (pAJ1181) and L1 stalk∆ (pAJ3605) in strain BY4741. D) Sucrose gradient sedimentation and northern blot analysis using oligo (AJO628) against a unique tag on rRNA expressed from WT (pAJ1181) or L1 stalk∆ (pAJ3605) rRNA. Total 25S and 18S rRNAs were detected using oligos AJO192 and AJO190, respectively.

### Pre-60S subunits without Rpl1 fail to recruit Mex67-Mtr2 heterodimer efficiently

The accumulation of Rpl25 and various shuttling biogenesis factors in the nucleus suggested that nascent 60S subunits lacking Rpl1 were defective in nuclear export. Possibly, nascent subunits lacking Rpl1 were unable to recruit factors involved in 60S export because of structural differences caused by loss of Rpl1. To identify such factors, we affinity purified nascent subunits and performed mass spectrometric proteomic analysis on them. After observing that Nmd3 can bind to large subunit particles lacking Rpl1 (Fig 2A-C), we decided to use C-terminal TAP-tagged Nmd3 as a bait for affinity purifying Pre-60S particles in *RPL1* repressed cells. As shown above, loss of Rpl1 from the pre-60S particle affected their nuclear export and accumulated particles in the nucleoplasm. For comparison, we affinity purified Nmd3-TAP particles from cells treated with LMB in an LMB-sensitive *CRM1-T539C* background (Grosshans et al. 2001), to mimic the nuclear accumulation of Rpl1-containing particles.

Spectral counts obtained from mass spectrometric analysis of the eluted samples were used to generate relative spectral abundance factor (RSAF) values as described previously (Sardana et al. 2015). We then generated ratios for RSAF values for each protein in the sample to that of Tif6 protein in the same sample and normalized values to the L1-expressed + LMB samples. Figure 4A summarizes results from two independent experiments, comparing the relative RSAF values for the 60S export factors Arx1, Bud20 and Mex67. While depletion of Rpl1 had no effect on the association of Arx1 or Bud20 with Nmd3-bound pre-60S particles, Mex67 was not detectable on these particles (Fig 4A). The loss of Mex67 from these particles was not due to reduced expression of Mex67 in the Rpl1-repressed cells (Fig 4C).

**Figure 4.**
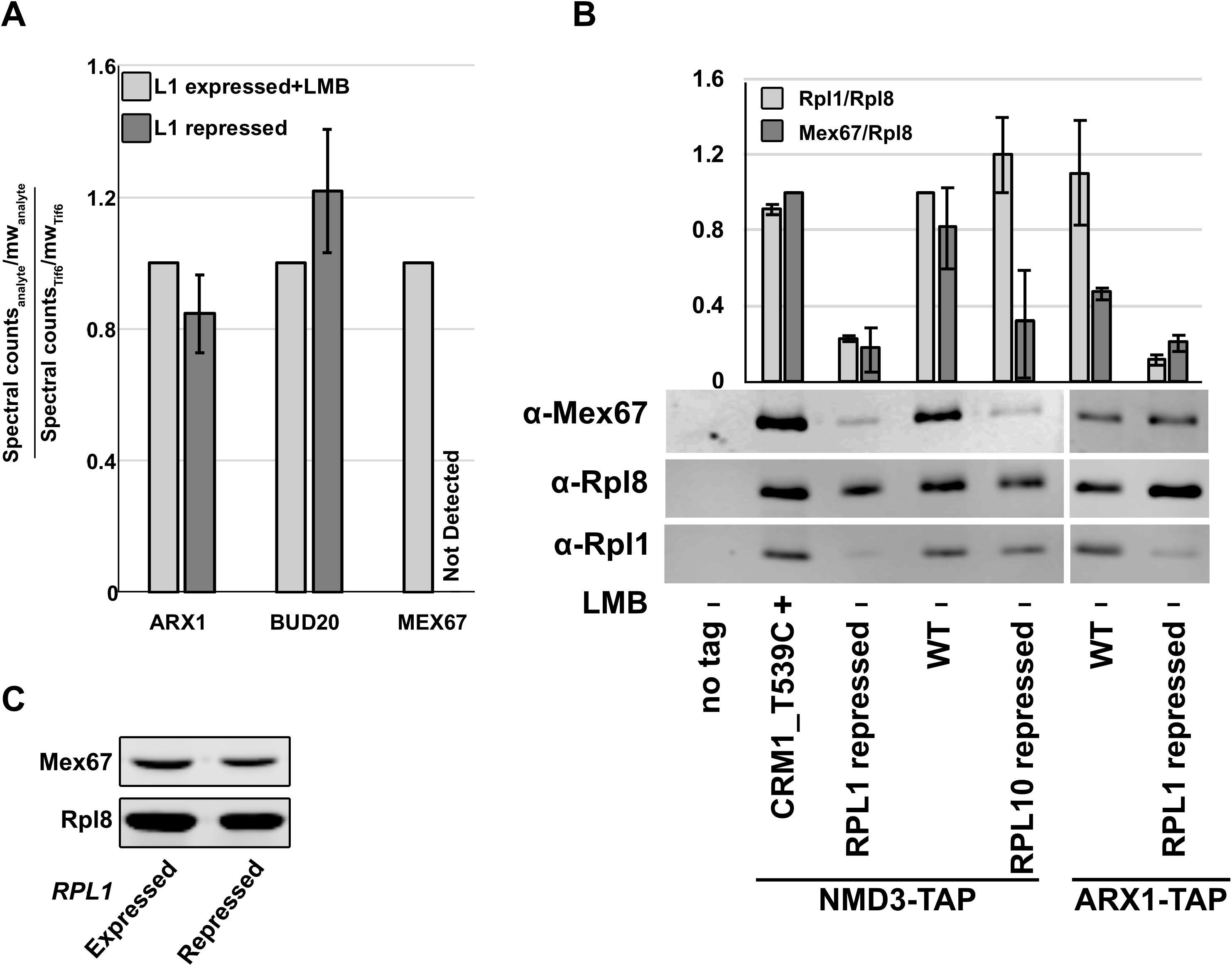
Nascent subunits lacking Rpl1 fail to recruit Mex67 efficiently. A) Nmd3-TAP was purified from *RPL1*-repressed cells and from LMB-treated cells. Spectral counts for Arx1, Bud20 and Mex67 were normalized to Tif6 levels in each sample. TAP purifications were from AJY4008 treated with LMB for 30 minutes and from AJY4009 in which *RPL1* was repressed for 1.5 hours. B) Western blots for Mex67, Rpl1 and Rpl8 in TAP purification samples from BY4741, AJY4008 treated with LMB for 30 minutes, AJY4009, AJY1874, AJY4001, AJY4012 and AJY4013 grown in galactose followed by 1.5h glucose treatment (lanes 1-7 respectively). Mex67:Rpl8 and Rpl1:Rpl8 were calculated for each sample. Mex67:Rpl8 ratio in each sample was normalized to that in the LMB sample (lane 2), and Rpl1:Rpl8 ratio in each sample was normalized to that in the WT NMD3-TAP samples (lane 4). C) Western blots for Mex67 and Rpl8 in extracts from KBM13 and KBM20 grown in galactose containing media for 48h and then diluted and grown in fresh glucose containing medium for 1.5h.

To corroborate the results from mass spec, we also analyzed both the Rpl1-repressed, and the Rpl1-containing and LMB-treated samples by SDS-PAGE and western blotting for Rpl8, Rpl1 and Mex67. For additional controls, we carried out mock TAP purification from untagged cells as well as Nmd3-TAP purification from wild-type and Rpl10-repressed cells, to trap particles after export and at a very late step in cytoplasmic maturation at which Mex67 would be expected to have already been released. Finally, as an additional control experiment, particles were purified using Arx1-TAP from WT or RPL1-repressed cells. Similar to the mass spec analysis, the amount of Mex67 co-precipitating with Nmd3-bound particles sharply decreased in *RPL1*-repressed cells compared to LMB-treated cells (Figure 4B, lanes 2 and 3), suggesting that pre-60S particles devoid of Rpl1 inefficiently recruited Mex67. In samples from WT cells, without any LMB treatment (lane 4), relative Mex67 levels were comparable to those in the LMB-treated sample. The RPL10-repressed sample also exhibited a sharp decrease in Mex67 levels (Figure 4B lane 5), as expected for a late-cytoplasmic particle. In Arx1-TAP samples too, less Mex67 was co-precipitated from RPL1-repressed cells compared to WT cells (Fig 4B lanes 6 and 7). However, the decrease was subtle compared to Nmd3-TAP particles perhaps because Arx1 binds pre-60S earlier than Nmd3, significantly and before Mex67 and hence a smaller population of Arx1 particles is bound to Mex67-containing particles.

### High copy expression of the Mex67-Mtr2 heterodimer specifically suppresses the export defect of Rpl1 repression

The low levels of nuclear export factor Mex67 associated with nascent 60S subunits purified from Rpl1-repressed cells suggested that Rpl1 may have a role in recruiting the Mex67-Mtr2 heterodimer to nuclear pre-60S. To test if the export block could be overcome, we co-overexpressed Mex67 and Mtr2 in *rpl1B*∆ cells with *RPL1A* under galactose inducible/glucose repressible promotor also expressing Rpl25-GFP. As shown in Figure 1, Rpl25-GFP accumulated in the nucleus upon repression of *RPL1A* but remained cytoplasmic under the same conditions in the WT cells (Figure 5A). However, co-overexpression of Mex67 and Mtr2 alleviated the nuclear export defect of nascent subunits, monitored by Rpl25-GFP localization. To test if the effect of Mex67-Mtr2 overexpression was specific to these export factors, we asked if over-expressing other 60S nuclear export factors, Arx1 and Bud20, could mitigate the nuclear export defect caused by Rpl1 loss. Overexpression of neither of these two proteins affected the nuclear localization of Rpl25, suggesting that the Mex67-Mtr2 binding is specifically affected upon Rpl1 loss. Although overexpression of Mex67 and Mtr2 suppressed the nuclear export defect of Rpl1 repression, co-overexpression of Mex67 and Mtr2 did not suppress the lethality caused by repression of Rpl1 (Figure 5B), as expected because Rpl1 is an essential ribosomal protein that interacts with E site ligands during the translation cycle.

**Figure 5.**
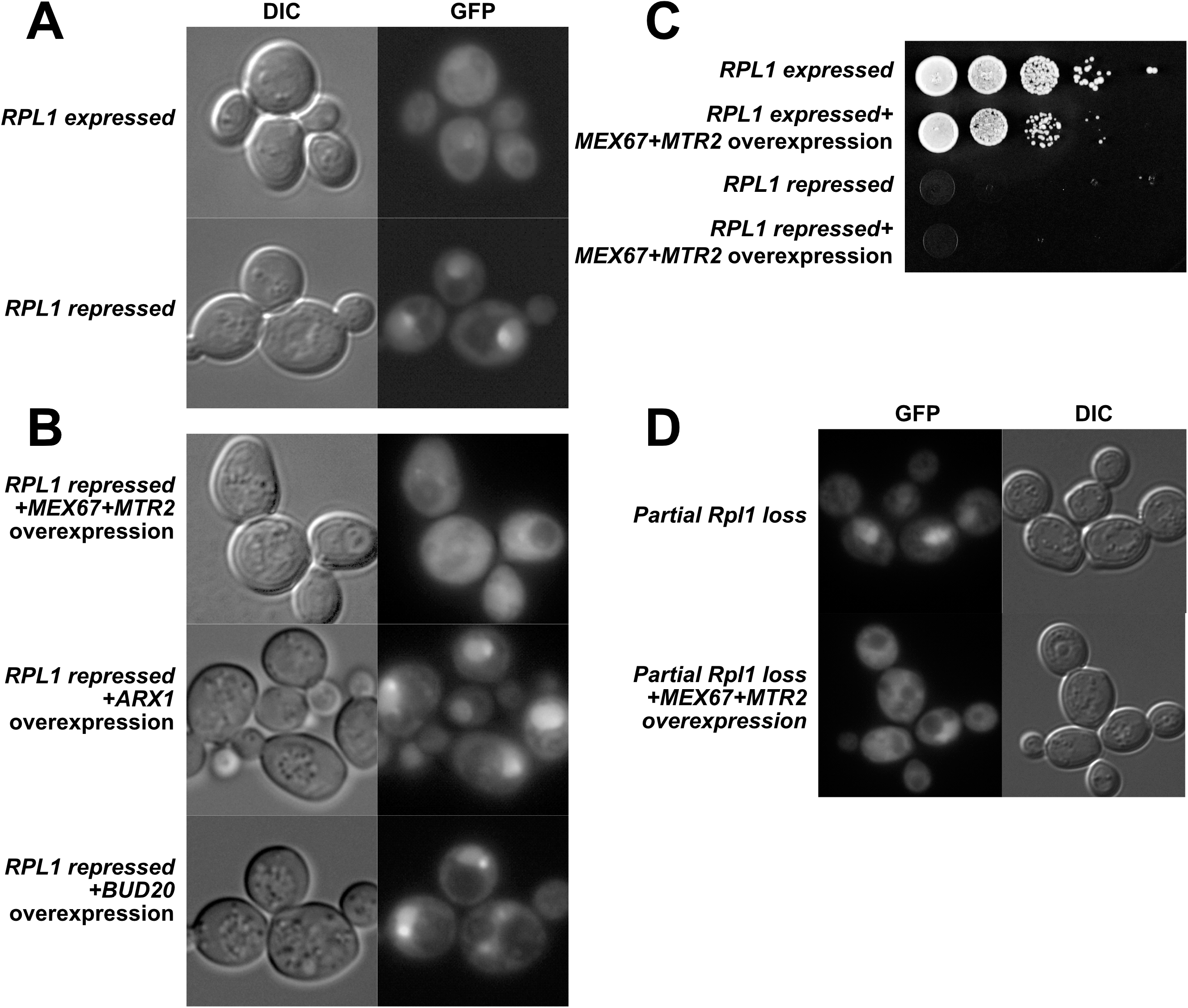
High copy expression of Mex67-Mtr2 heterodimer specifically suppresses the 60S export defect caused by Rpl1 loss. Rpl25-GFP viewed in A) WT (KBM13) transformed with *RPL25-GFP* (pAJ907) and empty vector (pRS426) and in *rpl1B*∆ *P*_*GAL-*_RPL1A (KBM20) with *RPL25-GFP* and empty vector or B) high copy *MEX67+MTR2* (pAJ3972), *ARX1* (pAJ4315) or *BUD20* (pAJ4316) and grown in Leu-Ura-media with galactose for 48h and then diluted 5-folds in glucose containing media and grown for 1.5 hours more. C) 10-fold serial dilutions of the KBM13 or KBM20 transformed with empty vector or *MEX67+MTR2* were spotted on glucose-containing selective media to repress *P*_*GAL*_:RPL1A in KBM20. D) Rpl25-GFP viewed in upper panel: *rpl1B*∆ (KBM17) transformed with *RPL25-GFP* (pAJ907) and empty vector (pRS426) and lower panel: KBM17 transformed with *RPL25-GFP* and *MEX67 MTR2* and grown in Leu-Ura-media with glucose for 48h and then diluted 5-fold in glucose-containing media and grown for 1.5 hours more.

## Discussion

Here, we have shown that the L1 stalk is needed for efficient export of pre-60S subunits from the nucleus. Although it was previously demonstrated that ribosomes lacking Rpl1 can engage in translation and therefore must be exported (McIntosh et al. 2011; Shi et al. 2017; Segev and Gerst 2018), those studies did not explore a role for Rpl1 in nuclear export. Considering that Rpl1 is essential in yeast and provides part of the binding site for the nuclear export adapter Nmd3, it is somewhat surprising that loss of Rpl1 does not have a greater impact on 60S export. Nmd3 is conserved from archaea to higher eukaryotes, indicating that it has a more fundamental role in ribosome maturation that predates evolution of the nuclear envelope. Whereas the euryarchaeal proteins are similar to eukaryotic Nmd3 and contain an eIF5A-like domain which interacts with Rpl1, the lower archaeal Nmd3 proteins lack this domain. Thus, the conserved function of Nmd3, which is likely to promote the loading of Rpl10 (uL16) (Zhou et al. 2019), is independent of Rpl1 binding. The interaction with Rpl1 appears to be a more recent evolutionary development and may not be essential for Nmd3 function.

### Quality control and the L1 stalk

Despite the essential nature of the L1 stalk and the expectation that quality control mechanisms monitor the nascent subunit for correct assembly, our work demonstrates that there is not a strict quality control pathway that assesses assembly of the L1 stalk. A similar observation was recently made for the RNA of internal transcribed spacer 2 (ITS2), which connects 5.8S and 25S RNA. The 232 nucleotides of ITS2 are normally removed by RNA processing in the nucleolus, prior to export. However, in mutants that are blocked for ITS2 processing, pre-60S subunits retaining ITS2 RNA are exported to the cytoplasm (Sarkar et al. 2017). These ITS2-containing subunits also engage with 40S subunits in translating ribosomes. However, they appear to induce a translational stress and are recognized by cytoplasmic quality control pathways involving 3’-RNA decay machinery and the RQC complex. In both cases, whether the defective ribosomes themselves are targeted for degradation and/or induce degradation of associated mRNAs remains to be resolved.

### What is the role of the L1 stalk in large subunit export?

The L1 stalk could impact export by one of a couple different but not mutually exclusive mechanisms. Because the L1 stalk is a highly flexible hydrophilic appendage, it might be unfavorable for passage through the NPC. Closing the L1 stalk by binding to Nmd3 would present a more compact structure to facilitate export. Similarly, expansion segment 27 (ES27) forms a long dynamic helix in the vicinity of the exit tunnel and is captured by the export factor Arx1, restraining its movement (Greber et al. 2016). Conceivably, tethering both the L1 stalk and ES27 could be mechanisms to facilitate export. In an attempt to ask if reducing the length of the L1 stalk could enhance export by eliminating a “floppy” RNA element, we made further truncations of the L1 stalk. However, we did not observe enhanced export of these larger L1 stalk truncations (data not shown).

Alternatively, the L1 stalk may be important for efficient recruitment of an export factor. The translocation of large cargo molecules through the nuclear pore complex requites multiple receptors (Ribbeck and Görlich 2002). Indeed, nuclear export of pre-60S subunits in yeast requires the export adapter Nmd3 (which recruits the receptor Crm1), the mRNA export receptor Mex67-Mtr2 and as well as Arx1 and Bud20 (Altvater et al. 2012; Bassler et al. 2012). Whereas the binding sites for Nmd3, Arx1 and Bud20 are well-established, the binding site for Mex67-Mtr2 has been enigmatic. Although the Mex67-Mtr2 duplex can bind 5S rRNA (Yao et al. 2007), *in vitro* UV-induced crosslinking of Mex67 reconstituted with pre-60S particles affinity purified with Yvh1, identified binding sites in the P stalk and 5.8S but not 5S rRNA (Sarkar et al. 2016). However, our recent structural analysis of pre-60S maturation (Zhou et al. 2019) shows that Yvh1 is recruited to the pre-60S only after Nog1 is released in the cytoplasm, a conclusion reached by others as well (Nerurkar et al. 2018). Thus, Yvh1 loads onto the pre-60S particle after export from the nucleus, and after the requirement for Mex67 in export. We suggest that in the Yvh1-bound particle, P stalk RNA is exposed due to the absence of either Mrt4 or the stalk protein P0, possibly offering a site for promiscuous binding by Mex67-Mtr2. UV-induced crosslinking of Mex67 to RNAs *in vivo* identified a wide distribution of crosslinking sites in 25S and 5.8S rRNA (Tuck and Tollervey 2013) with a strong hits in 5.8S, overlapping what was found *in vitro* (Sarkar et al. 2016). In neither of these crosslinking studies was Mex67 crosslinking to the L1 stalk observed. In addition, we did not detect interaction between Mex67 and the L1 stalk by yeast three hybrid (data not shown). Nevertheless, it is possible that Mex67 recruitment to the particle is enhanced by closure of the L1 stalk, by making a binding site in the vicinity of the L1 stalk, possibly 5.8S, more accessible.

After export to the cytoplasm, the pre-60S undergoes a series of maturation events culminating in the completion of the peptidyl transferase center and release of Nmd3 and Tif6. We previously observed Nmd3 bound to the L1 stalk in partially closed states (Malyutin et al. 2017) and suggested that the L1 stalk may be required for the release of Nmd3. However, the accumulation of Nmd3 in the nucleus in the absence of L1 expression argues against a requirement for L1 in the removal of Nmd3. Similarly, mutations in Nmd3 that are predicted to disrupt its interaction with L1 have only a very modest impact on growth, contrary to what would be expected if the L1-Nmd3 interaction were necessary for the release of Nmd3 (data not shown).

### L1 stalk mutants in translation

Rpl1 facilitates translation elongation assisting the release of E-site tRNAs and binding factors including eIF5A (Melnikov et al. 2016; Voorhees et al. 2009). Although the mechanism of translation is highly conserved, L1 is not essential in *E.coli* (Subramaniam and Dabbs 1980). It is essential in yeast but recent studies in both yeast and mammalian cells have detected L1 deficient ribosomes in actively translating pools (McIntosh et al. 2011; Shi et al. 2017). It has been suggested that yeast Rpl1-deficient ribosomes associated with polysomes are strongly discriminated against during translation initiation and a fraction of them is targeted for degradation (McIntosh et al. 2011). It has also been suggested that Rpl1 is required in “specialized ribosomes” for translating a specific subset of transcripts (Segev and Gerst 2018; Shi et al. 2017). Consistent with that, we show that ribosomes with truncated L1 stalk rRNA were able to engage in translation. However, the mutant ribosomes showed a strong bias towards lighter polysomes compared to wild-type ribosomes, possibly reflecting a general defect in elongation. Alternatively, ribosomes without an L1 stalk may support elongation at very low rates and induce more frequent stalling.

## Materials and Methods

### Strains plasmids and growth media

*S. cerevisiae* and plasmids used in this study are listed in Tables I and II. All cells were grown at 30°C in rich media (yeast extract and peptone) or synthetic dropout medium supplemented with 2% glucose or 1% galactose. Strains AJY3848, AJY3849, and AJY3850 were generated by genomic integration of *TIF6-GFP-HIS3MX, ARX1-GFP-HIS3MX* and *MRT4-GFP-HIS3MX*, amplified from AJY2909, AJY1948 and AJY3040, respectively, into KBM20. AJY4060 was generated by sporulation of KBM20 after mating with AJY1705. Strains AJY4001, AJY4008 and AJY4009 were generated by genomic integration of *NMD3-TAP-HIS3MX* amplified from AJY1874 into AJY3373, KBM13 and KBM20, respectively. AJY4012 and AJY4013 were generated by genomic integration of *ARX1-TAP-HIS3MX* amplified from AJY2491 into KBM13 and KBM20, respectively.

**Table I.**
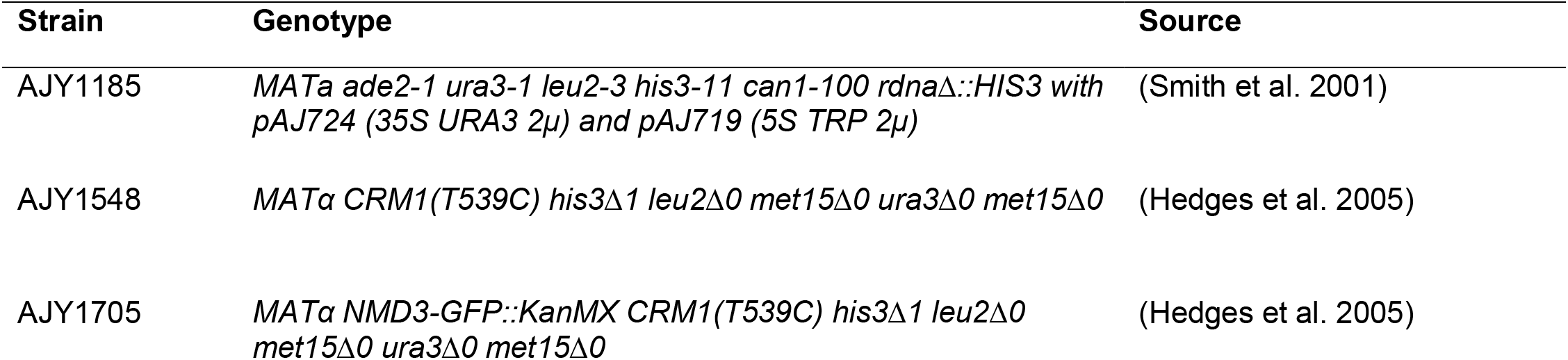
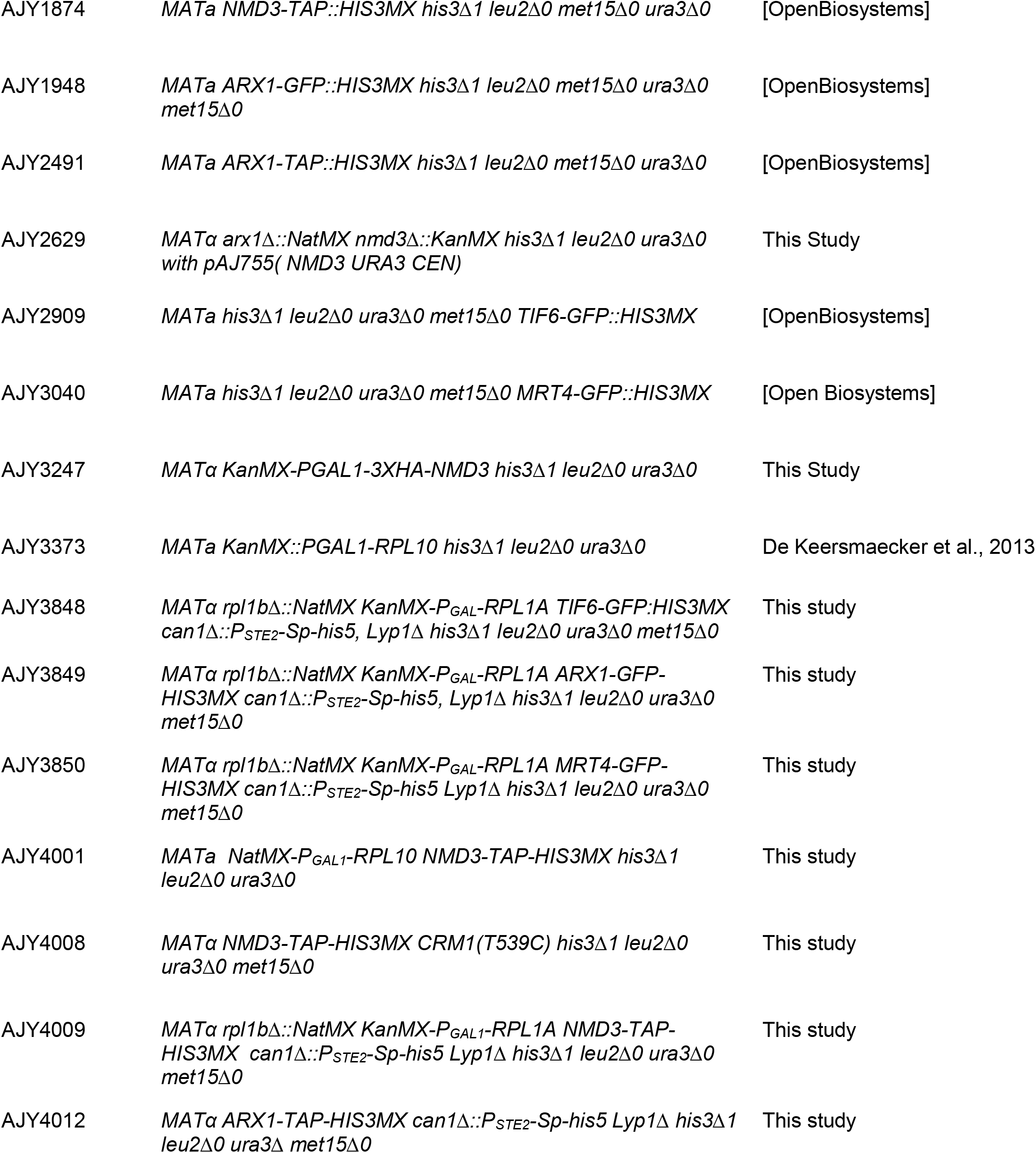
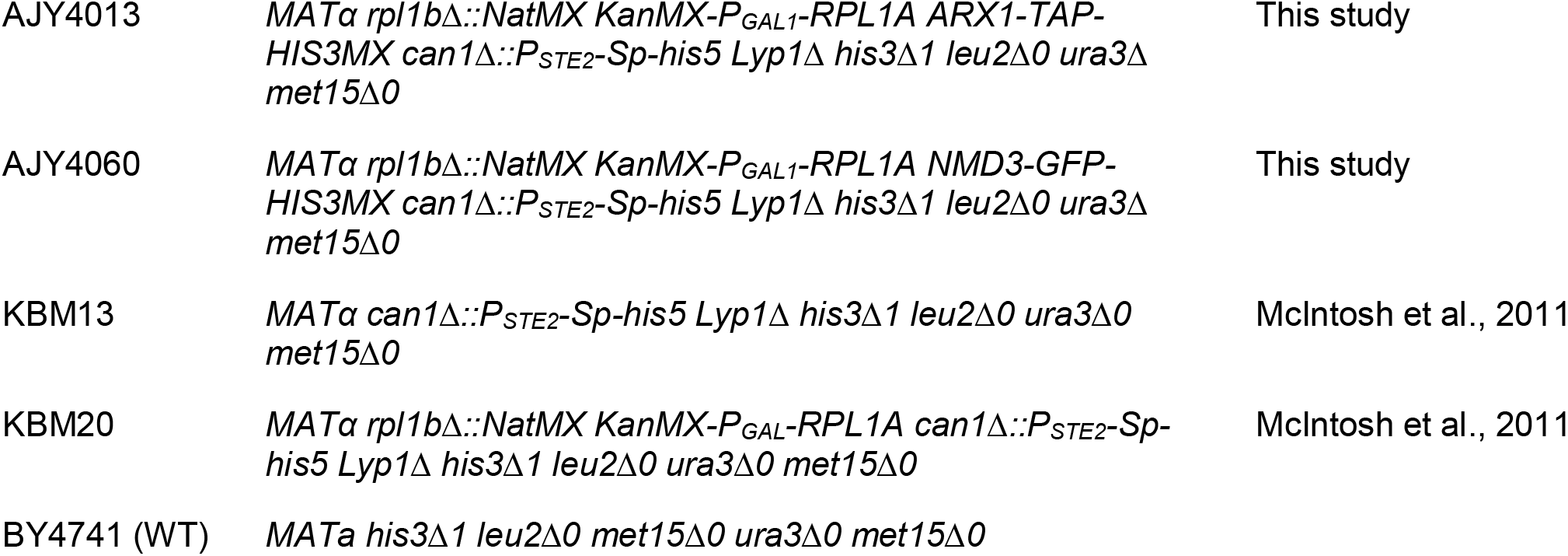
Strains used in this study

**Table II.**
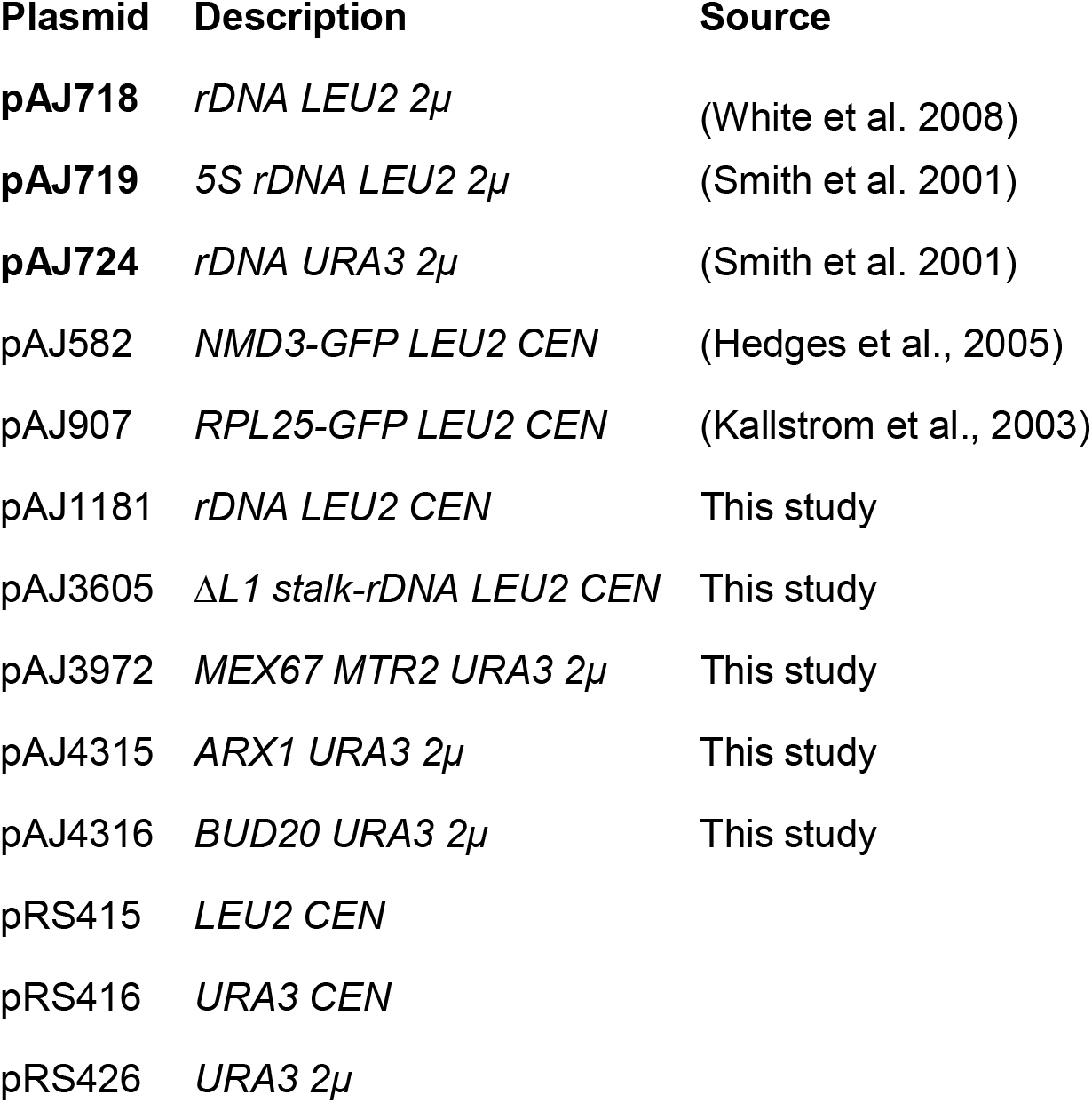
Plasmids used in this study

| Plasmid | Description | Source |
| --- | --- | --- |
| pAJ718 | *rDNA LEU2 2μ* | (White et al. 2008) |
| pAJ719 | *5S rDNA LEU2 2μ* | (Smith et al. 2001) |
| pAJ724 | *rDNA URA3 2μ* | (Smith et al. 2001) |
| pAJ582 | *NMD3-GFP LEU2 CEN* | (Hedges et al., 2005) |
| pAJ907 | *RPL25-GFP LEU2 CEN* | (Kallstrom et al., 2003) |
| pAJ1181 | *rDNA LEU2 CEN* | This study |
| pAJ3605 | *ΔL1 stalk-rDNA LEU2 CEN* | This study |
| pAJ3972 | *MEX67 MTR2 URA3 2μ* | This study |
| pAJ4315 | *ARX1 URA3 2μ* | This study |
| pAJ4316 | *BUD20 URA3 2μ* | This study |
| pRS415 | *LEU2 CEN* |  |
| pRS416 | *URA3 CEN* |  |
| pRS426 | *URA3 2μ* |  |

### Affinity purification of Nmd3-TAP and Arx1-TAP particles

Cultures of strains AJY1874, AJY4001, AJY4008, AJY4009, AJY4012 and AJY4013 were grown to OD_600_ of 0.3 in 500ml of YP media supplemented with 1% galactose. Glucose was added to a final 2% (w/v) concentration and cells were grown for one hour, harvested and cell pellets were frozen at −80°C. Cell pellets were washed and resuspended in 1.5 volumes of lysis buffer (20mM HEPES, pH 7.5, 10mM MgCl_2_,100mM KCl, 5mM β-mercaptoethanol, 1mM phenylmethylsulfonyl fluoride (PMSF), 1µM leupeptin, and 1µM pepstatin). Extracts were prepared by glass bead lysis and clarified by centrifugation at 4°C for 15 minutes at 18,000g. NP-40 was added to a final concentration of 0.15%(v/v) to the clarified extract which was then incubated with rabbit IgG (Sigma) coupled Dynabeads (Invitrogen) for 1h at 4°C. The Dynabeads were prepared as previously described(Oeffinger et al. 2007). Beads were then washed thrice with lysis buffer containing 0.15% NP-40 at 4°C for 5 minutes each time. The bound complexes were enzymatically eluted with tobacco etch virus protease in lysis buffer containing 0.15% NP-40 and 1mM Dithiothreitol for 90 minutes at 16°C.

### Polysome Analysis and western blots

Cultures of strains KBM13and KBM20were grown to OD_600_ of 0.3 in 150ml of YP media supplemented with 1% galactose. Glucose was added to a final 2% (w/v) concentration and cells were grown for two more hours. Cycloheximide (CHX) was added to a final concentration of 100µg/ml, cultures incubated for 10 minutes at 30°C to arrest translation and preserve polysomes and cells were harvested and frozen at −80°C. Cell pellets were washed and resuspended in 1.5 volumes of polysome lysis buffer (20mM HEPES, pH 7.5, 10mM MgCl_2_,100mM KCl, 100µg/ml CHX, 5mM β-mercaptoethanol, 1mM PMSF, 1µM leupeptin, and 1µM pepstatin). Extracts were prepared by glass bead lysis and clarified by centrifugation at 4°C for 15 minutes at 18,000g. 9 A_260_ units of clarified extract were loaded onto 7-47% sucrose gradients prepared in polysome lysis buffer and centrifuged for 2.5 hours at 40,000 rpm in a Beckman SW40 rotor. Gradients were fractionated using an ISCO Model 640 fractionator into 600µl fractions with continuous monitoring at 254nm. 1.2 ml 100% ethanol was added to each fraction, vortexed and stored at −20°C overnight. Fractions were centrifuged at 4°C for 15 minutes at 18,000g and pellets were dissolved in 1X Laemmli buffer and heated at 99°C for 3 mins. Proteins were separated on 6-15% gradient SDS-PAGE gels, transferred to nitrocellulose membrane and subjected to western blot analysis using anti-Nmd3, anti-Rpl8 (K.-Y. Lo) and anti-Rpl1 (J. Warner) antisera.

### Sucrose gradient sedimentation and northern Blot Analyses

Saturated cultures of BY4741 transformed with pAJ1181 or pAJ3605 were diluted to OD_600_ of 0.1 in SD Leu^−^ and grown to mid log phase. Cell cultures were treated with CHX at 50µg/ml for 10 mins at 30°C to inhibit translation and then cells were harvested and stored at −80°C. Cells were washed once and then resuspended in 1.5-2 volumes lysis buffer (50 mM Tris-HCl pH 7.5, 100 mM KCl, 5 mM MgCl2, 50 µg/mL CHX, 1 mM PMSF, benzamidine, and 1 µM each of leupeptin and pepstatin). Extracts were prepared by glass bead lysis and clarified by centrifugation at 4°C for 15 minutes at 18,000g. 9 A_260_ units of clarified extract were applied to sucrose density gradients and fractionated as described above. 60µl of 20% SDS, 60µl of 3M Sodium acetate pH5.2 and 1.3ml 100% ethanol were added to each sample and nucleic acids were precipitated overnight at −20°C. The precipitate was pelleted by centrifugation for 30mins at 18000 rpm and 4°C. RNA pellets were washed with 70% ethanol and air dried. Pellets from each fraction were dissolved in 50µl RNAse free water. Total RNA from one A_260_ unit of clarified lysate was extracted similarly and dissolved in 50µl RNAse free water. 10ul RNA samples from lysate and from sucrose gradient fractions 1-19 were vacuum dried and dissolved in 10 ul RNA sample loading buffer (Invitrogen AM8552). RNAs were resolved by electrophoresis through 1.2%-agarose MOPS 6% formaldehyde gel for 4 h at 50 volts. Northern blotting was performed as previously described (Li et al. 2009) using the oligos AJO190, AJO192 and AJO628 (Table III), and signal was detected by phosphoimaging on a GE Typhoon FLA9500.

**Table III.**
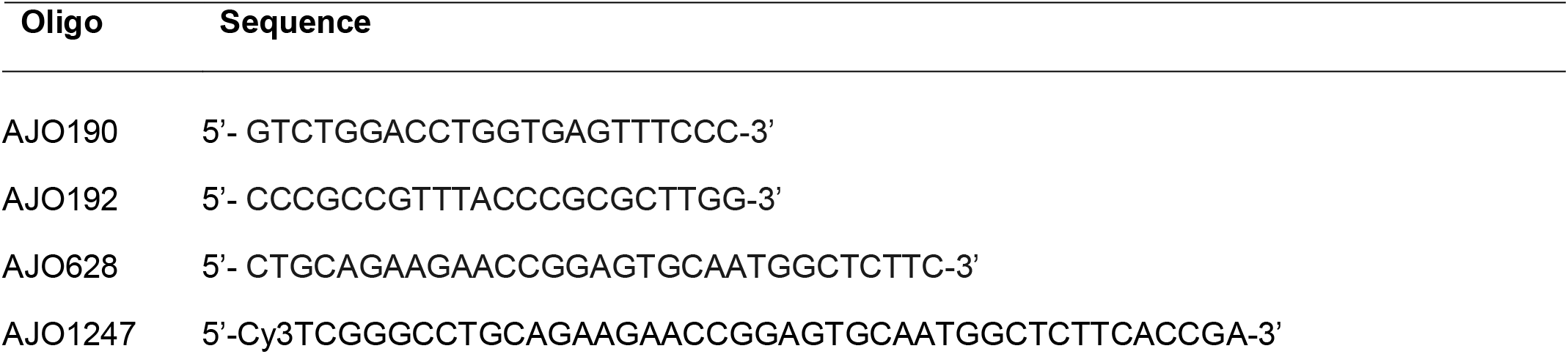
Oligonucleotides used in this study

### Microscopy

For direct fluorescence experiments, cells were grown in selective medium (Leu^−^ or His^−^) supplemented with 1% galactose for 48 hours, then diluted 4-fold in medium containing 2% glucose and grown for 60-90 minutes to repress the expression of *RPL1A*. Images were captured using a Nikon E800 microscope fitted with a 100× Plan Apo objective and a Photometrics CoolSNAP ES camera controlled by NIS-Elements software. For fluorescence *in situ* hybridization experiments BY4741 cells transformed with pAJ1181 or pAJ3605 were grown to saturation in Leu^−^ medium with 2% glucose, then diluted 5-fold in fresh Leu^−^ glucose medium and continued to grow for 60 minutes. Formaldehyde was added to a final concentration of 4.5% to the cell cultures and cells were fixed by agitating gently at room temperature for 30 minutes. Fixed cells were pelleted and washed twice with KSorb buffer (1.2M sorbitol, 0.1M potassium phosphate buffer 7.0). Cell pellets resuspended in 200ul KSorb, were treated with 50µg/ml Zymolyase T20 for 15 minutes at 37°C in presence of 20mM Vanadyl Ribonucleoside complex (VRC), 28mM ꞵ-mercaptoethanol and 1mM PMSF. Cells were gently pelleted and washed with ice cold KSorb buffer thrice and resuspended in 100µl Ksorb buffer. 35µl cell suspension was applied to the wells of Teflon coated Immunofluorescence slides (Polysciences Inc, No. 18357) pre-coated with Poly-lysine. Slides were incubated in a moist chamber at room temperature for 10 mins, then excess cells were gently aspirated, and the slides were stored in 70% ethanol at −20°C. Cells were rehydrated by washing twice with 2X SSC (300mM NaCl, 30mM Sodium Citrate pH 7.0) and then incubated in 40µl Prehybridization solution (10% Dextran sulfate, 50% deionized formamide, 1X Denhardt’s, 2mM VRC and 4X SSC, 0.2% BSA, 25µg yeast tRNA and 500µg/ml ssDNA) for 1h at 72°C in a moist chamber. Excess solution was removed by aspiration and replaced with 40µl of Hybridization solution (Prehybridization solution containing 1µM Cy3 labelled oligo AJO1247) in each well. The slide was incubated in a moist chamber at 72°C for 1h followed by overnight incubation at 37°C. Next day, the wells were washed with 2X SSC and then 1X SSC containing 0.1% NP-40 for 30 minutes each. Cells were incubated for 2 minutes with 1µg/ml DAPI, washed twice with PBS and mounted in Aqua-Poly/Mount (Polysciences, Inc). Images were captured as described above.

## Author Contributions

S.M. and A.W.J. designed the study. S.M. designed and performed the experiments. J.B. performed the northern blot analysis. S.M. and A.W.J. interpreted the results and wrote the manuscript.

## Acknowledgements

We thank Dr. J. R. Warner (Albert Einstein College of Medicine, New York) for his generous gift of yeast strains and anti-Rpl1 antibodies, Dr. K.-Y. Lo for anti-Rpl8 antibody and members of the Johnson lab for helpful discussions. This work was supported by NIH Grants GM53655 and GM127127 (to A.W.J.).

